# Connectivity and phenotype of vasopressin 1a receptor cells in the lateral septum

**DOI:** 10.1101/2025.10.26.683946

**Authors:** Delenn Hartswick, Arkar Zaw, Nicholas Schappaugh, Caitlin N. Friesen, Geert J. de Vries, Aras Petrulis

## Abstract

The neuropeptide arginine-vasopressin (AVP) regulates sexually-differentiated social behaviors, including sexual behavior, aggression, and social communication. Much of AVP’s effect on social behavior is mediated by the most widely-expressed AVP receptor in the central nervous system: the vasopressin 1a receptor (V1aR). Dense expression of V1aR is found in males and females in the lateral septum (LS), which also receives heavy input from sexually-differentiated populations of AVP cells in the extended amygdala. In order to access and interrogate the structure and function of V1aR cells, we developed and validated a V1aR-Cre knockin mouse line (Avpr1a-P2A-iCre). V1aR-Cre mice did not differ from their Cre-negative littermates in their health, sensorimotor function, anxiety/motivational behavioral responses, or in their V1aR binding levels in the LS. The distribution of Cre in the brain, as assessed by crosses with Cre-reporter mouse lines and *in situ* hybridization (ISH), was highly similar to patterns of V1aR binding and demonstrated strong colocalization between Cre and V1aR expression within LS cells. We used this V1aR-Cre mouse to identify the inputs and outputs of V1aR+ cells in the LS, via a monosynaptic retrograde rabies virus as well as anterograde synaptophysin-mRuby AAV tracing approaches. Monosynaptic inputs to V1aR LS cells were observed from the medial preoptic area (MPA), lateral hypothalamus (LH), hippocampus (mostly ventral), medial septum, diagonal band of Broca (DBB), and the supramammillary nucleus (SuM). Synaptophysin labeling revealed outputs to some of these same structures (MS, LH, DBB, SuM) as well as parts of the lateral preoptic area (LPO), anterior hypothalamic area (AHA), and ventral pallidum. Notably, most structures (except for hippocampus) bidirectionally connected to V1aR LS cells also contain populations of V1aR+ cells and are targets for BNST/MeA AVP cells, suggesting an integrated AVP/V1aR circuit. Finally, using ISH, we measured levels of V1aR mRNA expression in subregions of the LS and colocalization of V1aR with oxytocin receptor (OTR) mRNA, which also has a high affinity for AVP, within the LS. We found similar high percentages of cells containing V1aR+ puncta across dorsal and intermediate LS in both males and females. In contrast, the ventral LS contained fewer V1aR+ cells in both males and females. The highest level of co-expression of V1aR and OTR was found in the intermediate LS in both sexes, suggesting the possible location of functional interactions between AVP and oxytocin within the LS. We also sought to identify other phenotypic aspects of V1aR cells using ISH and confirmed that all LS V1aR cells are GABAergic and that most also express corticotropin-releasing hormone receptor 2, suggesting a substrate by which AVP could interact with stress-related systems in the LS. Our characterized V1aR-Cre mouse will be a valuable resource for understanding the role of AVP/V1aR in behavioral and physiological systems. Using this mouse, we have characterized the connectional architecture of V1aR cells in the LS and revealed an interlocking set of structures that may be the substrate through which AVP regulates social and emotional behavior.

## Introduction

The neuropeptide arginine vasopressin (AVP) is a highly conserved signaling molecule that regulates many aspects of social and emotional behavior across vertebrate species, including affiliation, aggression, parental care, and social recognition (Caldwell, 2017; Goodson and Bass, 2001; Insel, 2010; Kelly and Goodson, 2014; Rigney et al., 2022). In humans and other mammals, AVP influences complex aspects of social cognition and bonding (Caldwell et al., 2008; Zink et al. 2011), highlighting its important role in connecting social cues with emotional and motivational states (De Vries and Panzica, 2006; Newman, 1999). AVP is produced mainly in three hypothalamic nuclei: the paraventricular (PVN), supraoptic (SON), and suprachiasmatic (SCN) nuclei. These cells regulate essential physiological functions such as water and salt balance, blood pressure, and circadian rhythms (De Vries and Miller, 1999; Grinevich and Ludwig, 2021; Rohr et al., 2021). In addition to these classical regions, AVP is also expressed in parts of the extended amygdala, including the bed nucleus of the stria terminalis (BNST) and medial amygdala (MeA) (De Vries and Panzica, 2006). These regions are part of the social behavior network, a conserved set of brain areas that coordinate social behaviors across species (Newman, 1999; O’Connell and Hofmann, 2011). Notably, AVP expression in the BNST and MeA is sexually differentiated, with males exhibiting greater AVP cell numbers and denser AVP fiber innervation to projection targets than females (De Vries, 2008; De Vries and Panzica, 2006; Rigney et al., 2023; Rood et al., 2013). These steroid-dependent AVP populations play crucial roles in sex-specific regulation of social and emotional behaviors, including social investigation, aggression, and anxiety (Kelley and Goodson, 2013; Rigney et al., 2019; Rigney et al., 2024; Veenema et al., 2013; Whylings et al., 2021).

The vasopressin 1a receptor (V1aR) is the main mediator of AVP’s central nervous system effects (Koshimizu et al., 2012) on social interaction and interest (Patel et al., 2022; Rigney et al., 2024), social recognition (Bielsky et al., 2005; Landgraf et al., 1995), aggression and territorial behavior (Albers, 2012; Gobrogge et al., 2009), parental care (Bayerl and Bosch, 2019; Wang et al., 1994), and play behavior (Bredewold et al., 2014; Veenema et al., 2013). Tracing studies have revealed that the lateral septum (LS) is a major target of BNST/MeA AVP innervation (Caffé et al., 1987; De Vries and Buijs, 1983; Rigney et al., 2023) and is also rich in V1aR, identifying it as a key neural substrate for the behavioral actions of AVP (Besnard & Leroy, 2022; Bielsky et al., 2005; Bredewold and Veenema, 2019). Experimental manipulations of V1aR activity in the LS (i.e. pharmacological, genetic, and viral approaches) alter affiliative behavior, anxiety-like responses, and social recognition and approach, often in sex-dependent ways (Rigney et al., 2023; Veenema et al., 2013). For example, agonism of V1aR in the LS increased social play in male rats but decreased it in female rats (Bredewold et al., 2014; Veenema et al., 2013). Similarly, antagonism of V1aR in the LS, but not the MeA, in male mice impaired social recognition while AVP stimulation here potentiated social recognition (Bielsky et al., 2005). In addition, V1aR cells in the LS can also influence anxiety-like behavior, often in sex-specific ways. For example, male, but not female, V1aR knockout mice show reduced anxiety-like behavior (Bielsky et al., 2004, Bielsky et al., 2005), an effect that can be rescued by viral expression of V1aR in the LS (Bielsky et al., 2005). Recently, we have demonstrated that excitation of BNST AVP terminals in the LS increases anxiety-like behavior in a V1aR dependent manner in male, but not female mice (Rigney et al., 2024).

Despite the importance of LS V1aR in social and emotional processes, the circuit organization of V1aR-expressing neurons in the LS and their cellular identity remain poorly understood. Consequently, to fill these gaps in knowledge, we developed and validated a *Avpr1a-iCre* driver mouse line and used it to identify the circuit-level organization of LS V1aR cells in males and females using viral-vector anterograde and retrograde tracing approaches. We also used *in situ* hybridization (ISH) to better define the cellular identity of these cells by examining *Avpr1a* mRNA co-expression with markers of glutamatergic and GABAergic neurotransmission, neuropeptides, and receptor/trophic systems known to be prominently expressed in the LS. By characterizing LS V1aR cells and their inputs and outputs in both sexes, we can further our understanding of how AVP influences the brain and behavior.

## Materials and Methods

### Animals

Both adult male and female Avpr1a-P2A-iCre (*Avpr1a^Cre+^*) knock-in mice and their Cre-negative (*Avpr1a^Cre-^*) littermates (as controls) were used for behavioral testing, health assessment, and neural tracing. Heterozygous *Avpr1a^Cre+^* mice were crossed with ZsGreen (Ai6; B6.Cg-*Gt(ROSA)26Sortm6(CAG-ZsGreen1)Hze*/J; Jackson Laboratory #007906) or tdTomato (Ai14; B6.Cg-*Gt(ROSA)26Sortm14(CAG-tdTomato)Hze*/J; Jackson Laboratory #007914) cre-reporter mouse lines to identify Cre-dependent fluorophore expression linked to *Avpr1a* gene expression (*Avpr1a^Cre+/ZsGreen^, Avpr1a^Cre+/tdTom^*). Adult male and female wild-type C57BL/6J (Jackson Laboratory #000664) mice were used to determine colocalization of native expression of *Avpr1a* mRNA and other cellular markers as well as for baseline autoradiography experiments. All mice were kept on a 12:12 reverse light cycle at 22°C and single-housed (Allentown NexGen cages) with corn bedding (Bed-O-Cobb) with *ad libitum* access to food (5001 Rodent Diet) and water unless otherwise mentioned. All animal procedures were performed in accordance with the Georgia State University Animal Care and Use Committee regulations and the National Institutes of Health Guide for the Care and Use of Laboratory Animals.

### Surgical procedures

All surgeries were conducted under 1.5-3% isoflurane gas anesthesia in 100% oxygen. Carprofen (3mg/kg) was administered subcutaneously before surgery and carprofen-loaded medigels (3mg/2oz Medigel Sucralose, ClearH2O, Westbrook, ME, USA) were placed in cages for the three days following surgery for pain management. Mice were placed in a stereotaxic frame (David Kopf Instruments, Tujunga, CA, USA) with ear and incisor bars with bregma and lambda leveled on the dorsal-ventral axis. Once the head was secured and leveled, a midline scalp incision was made, and a hand-operated drill was used to drill a hole in the skull, exposing dura. Coordinates used were in relation to bregma in mm; lateral septum (LS) coordinates: DV: -3.25, AP: +0.35, ML: +/-0.5 (males), 0.45 (females). Incisions were closed with wound clips which were removed within two weeks.

### Generation and genotyping of *Avpr1a^Cre+^* knock-in mice

We generated Avpr1a-P2A-iCre (*Avpr1a^Cre+^*) knock-in mice using homologous recombination in embryonic stem (ES) cells. A knock-in targeting vector was constructed to insert a P2A-iCre cassette immediately upstream of the endogenous *Avpr1a* stop codon in exon 2. The vector contained ∼2 kb 5′ and 3′ homology arms flanking the insertion site, as well as a neomycin resistance (Neo) cassette flanked by FRT sites for positive selection. The P2A sequence encoded a self-cleaving peptide, enabling bicistronic expression of the vasopressin 1a receptor (V1aR) and a codon-optimized Cre recombinase (iCre) from a single transcript.

The targeting vector was electroporated into iTL BF1 hybrid embryonic stem (ES) cells (C57BL/6N × 129/Sv, expressing Flp recombinase; Ingenious Targeting Laboratory, Ronkonkoma, NY). Following G418 selection, resistant ES colonies were picked, expanded, and screened for homologous recombination by PCR using primers spanning the short homology arm (A2) and the Neo cassette (ivNeoN3). Correctly targeted clones yielded the expected 2.78 kb amplicon, whereas wild-type DNA produced no product. Positive clones were further validated by PCR with SQ1 and FN2A primers, confirming cassette retention with a 1.93 kb product. Sequence analysis across the 5′ junction with primer iCreERT2 SC1 verified proper insertion of the P2A-iCre cassette without disrupting the coding frame. Alignment to the reference *Avpr1a* transcript (Ensembl ENSMUST00000020323.6) confirmed precise integration, with only minor non-coding base-call ambiguities.

Validated iTL BF1 (C57BL/6 FLP) ES clones were microinjected into BALB/c blastocysts to generate chimeric animals. High-percentage black coat color chimeras were bred with C57BL/6N wild-type mice to establish germline transmission and obtain Neo-deleted offspring. Tail DNA from black-coated pups was analyzed to confirm excision of the FRT-flanked Neo cassette, which left a single FRT “scar” at the targeted locus. Verified Neo-excised animals were subsequently backcrossed to C57BL/6J wild-type mice to remove the Flp transgene. Offspring were genotyped by PCR to distinguish wild-type and P2A-iCre alleles. For detection of the iCre knock-in allele, the following primers were used: forward 5′-TCCTGGGCATTGCCTACAAC-3′ and reverse 5′-CTTCACTCTGATTCTGGCAATTTCG-3′, with TaqMan probe 5′-ACCCTGCTGCGCATTG-3′. Heterozygous *Avpr1a^Cre+^* mice were maintained by backcrossing to C57BL/6J, while Cre-negative (*Avpr1a^Cre-^*) littermates were used as wild-type controls.

### Health and behavioral assessment of *Avpr1a^Cre+^* knock-in mice

To confirm that the presence of the iCre knock-in did not significantly impact health or behavior of the *Avpr1a^Cre^* strain, we tested N2 generation adult male and female *Avpr1a^Cre+^* and their *Avpr1a^Cre-^* littermates (n=10 per sex/genotype; 3 cohorts) on a battery of health and behavior assessments (Figure 1). Before each test/assessment, subjects were brought into the testing suite and given at least 30 minutes to acclimate before testing began. All testing and assessment, except for the elevated plus maze (EPM), which was conducted under white light, was performed under red light during the lights-off portion of the subject’s light cycle.

**Figure 1.**
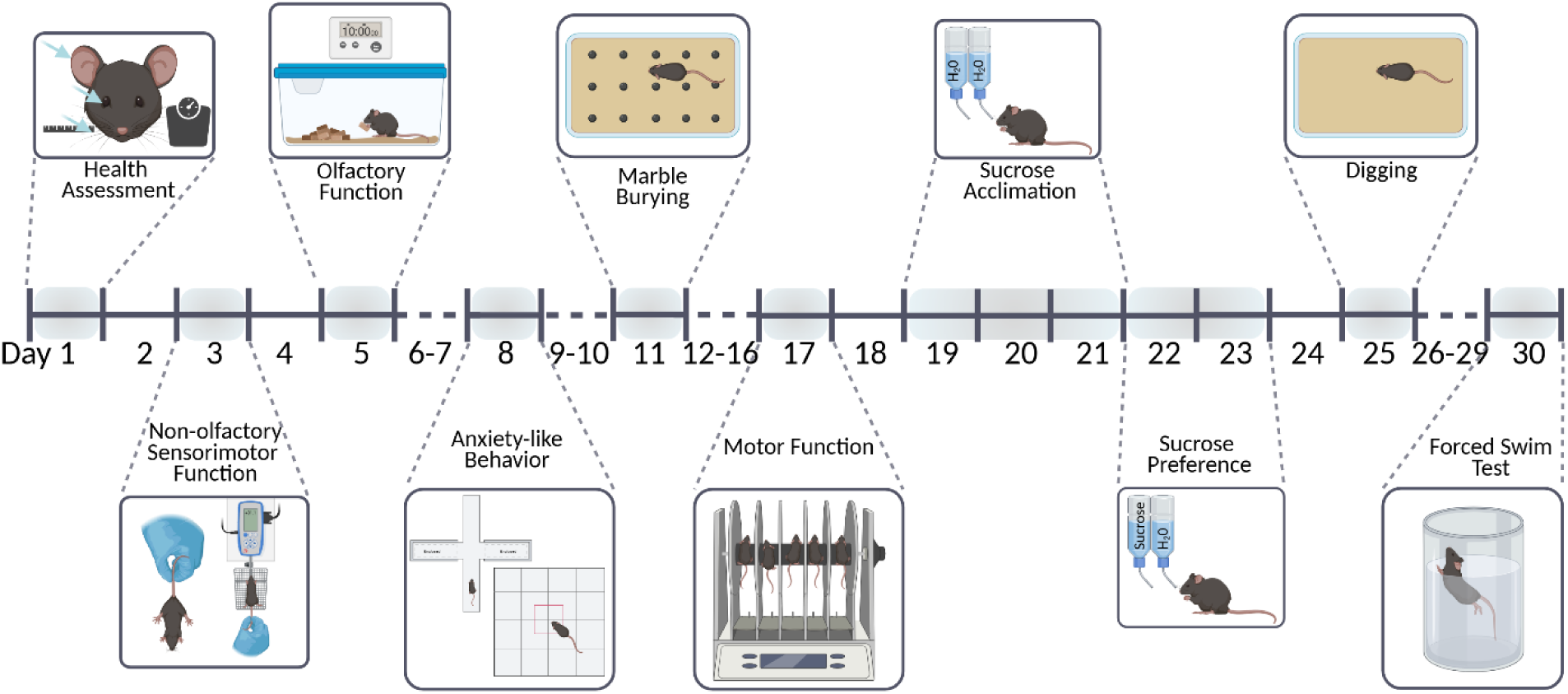
Timeline of health and behavioral assessments utilized to validate the Avpr1a-P2A-iCre knock-in strain. Assessments are described in detail in the methods section.

### Health assessment

Assessments were all done on the same day in the order listed. *Vibrissae*. Each subject’s whiskers were assessed for appropriate size and length, assigning them scores of 0 for whiskers being present and unbarbered, and a score of 4 for animals lacking any whiskers entirely. Intermediate lengths of whiskers at various levels of barbering were given scores of 1, 2, and 3 depending on the severity. *Eyes*. Subjects were checked to ensure they possess healthy eyes without any abnormalities in shape, size, or functionality. Subjects were given scores of 0 for both eyes being healthy, a score of 1 for any abnormalities in a single eye, or a score of 2 for abnormalities present in both eyes. *Ears*. The appearance of the pinna was assessed for their health and shape in subjects, independent of ear-punch for identification and genotyping. Normal and healthy ears were given a score of 0, any malformations of absence in a singular pinna resulted in a score of 1, and a score of 2 indicated such problems were present in both ear pinna. *Muscle Tone.* Subjects were put into the hands of the experimenter and allowed to climb their fingers to assess the normal functionality of muscles necessary for movement and strength. Subjects were given a score of 0 for normal muscle performance, a score of 1 if they had difficulty in climbing and holding onto the experimenter’s fingers, or a score of 2 if they showed significant muscular dysfunction such that normal movement was not possible. *Body Weight*. Subjects were lightly anesthetized via brief exposure to 4% isoflurane in 100% oxygen in an induction chamber, and then placed onto a scale to measure their body weight (grams). *Body Length*. Following body weight assessment, the anesthetized subjects were laid out onto a flat surface, and then a ruler was used to measure length (cm) from the tip of the nose to the base of the tail. Following this final assessment, subjects were returned to their cages to recover from anesthesia.

### Behavioral assessment

#### Motor function

Subjects were placed in a rotarod apparatus (Ugo Basile Mouse RotaRod, model 47650) and then the rods were rotated at 5 RPM, increasing at a constant rate to 40 RPM over the five-minute testing period. The latency to fall off the rod or inability to remain on top of the spinning rod for two subsequent rotations was measured. Subjects that maintained their position for the entire test were scored at maximum latency (300 s). Mice were immediately returned to their cage at the end of each session. Subjects were tested three times with at least 5 min between sessions, and their results were averaged across all three sessions.

#### Non-olfactory sensorimotor function

Sensorimotor function tests were all done on the same day in the order listed. *Visual placing.* This test was used to detect visual impairment across three trials. During each trial, subjects were held by the tail and slowly lowered toward the edge of a table, and a score of 1 was assigned if the subject extended its forepaws or 0 if it did not. The subject was determined to have passed the visual test if it had a cumulative score of at least 2 across the three trials.

#### Contact placing

This test was conducted to detect deficits in the sensory function of vibrissae across three trials. On each trial, a mechanical pencil was gently passed along one side of the subject’s head to touch the edge of its vibrissae. The side of the head was randomized across trials. The reaction to touch was evaluated and scored on a scale of 0-3 for each trial. For all subjects where vibrissae were present (a score of 0 was assigned if no vibrissae were present), a score of 1 was assigned if there was no response, 2 was assigned for a moderate response (e.g., small twitch), and 3 was assigned for a maximal response (e.g., head turn). The average score across three trials was used for analysis. *Limb clasping.* This was assessed for each subject by suspending the mouse by its tail approximately 30 cm above a table for 5 seconds across three trials. A score of 0 was assigned if all limbs were clasped against the body, a 1 was assigned if one limb one extended, a 2 was assigned if two to three limbs were extended, and a 3 was assigned if all limbs were extended. The limb clasping score was summed across all three trials. *Auditory function.* Hearing function was assessed by sounding a clicker 15 cm behind the subject’s head across three trials. A score of 1 was assigned if the subject flinched or turned its head, and a 0 was assigned if the subject did not respond. The subject was determined to have passed the test if it had a cumulative score of at least 2 across three trials. *Grip strength test.* Each subject’s forelimb grip strength was measured using a grip strength meter (Ugo Basile, #47200) across three trials. Each subject was placed on the pull bars, gently pulled, and the peak force was recorded by the meter. The average peak force across three trials was used for analysis.

#### Olfactory function

Subjects and their food and water were removed from their cage, and ten cereal pieces (Froot Loops; approximately 2g) were placed in a randomly chosen corner and covered with bedding. Mice were returned to their cage with the top covered with a perforated acrylic panel. The latency to find the cereal (when the mouse exposed buried cereal) was recorded. If the subject did not find the food over the ten-minute trial, it was assigned a 600-second latency score.

#### Anxiety-like behavior

To test for any changes in anxiety-like behavior caused by iCre insertion, we tested subjects on the elevated plus maze and in an open-field environment (Billah et al., 2019; Weiss et al., 2000). *Elevated Plus Maze*. The elevated plus maze (EPM) consisted of two open arms (80 cm x 5 cm) and two closed arms (80 cm x 5 cm x 15 cm) crossed perpendicularly and raised 51 cm above the floor. Subjects were placed into the center of the intersection and were allowed to habituate to the apparatus for 1 minute. The subject’s activity was then video recorded from above for an additional 5 minutes. Time spent in open and closed arms was determined by manual scoring from video recordings. *Open Field Test*. Subjects were placed in a 43 cm long x 43 wide x 30 cm tall open field chamber (Med Associates) for 10 minutes. The amount of time spent along the edges of the box and the time spent in the center area were automatically recorded (Activity Monitor software, Med Associates).

#### Sucrose preference

To assess whether iCre insertion altered hedonic processing, we tested subjects on their preference for consuming a palatable substance (Liu et al., 2015, Primo et al., 2023). During the three-day acclimation period, two water bottles were placed in each subject’s home cage. On the day of the sucrose preference test, subjects’ original water bottles were replaced with pre-weighed bottles: one containing a 2.5% sucrose solution in tap water, and the other containing tap water alone. Subjects had access to both bottles for the next 20 hours. The amount of liquid consumed from each bottle was measured after the bottles were removed. As a control, a water bottle was placed in a nearby empty cage and was monitored for the same period; this showed negligible (<1 mL) loss of solution. Preference for sucrose was calculated as the percentage of sucrose solution consumed divided by the total amount of liquid consumed (Sucrose/(Sucrose+Water) * 100%).

#### Marble burying and digging

To test whether iCre insertion altered repetitive behavior, we assessed each subject’s burying response to novel objects (De Brouwer et al., 2018, Dixit et al., 2020) as well as to a novel bedding condition (Angoa-Perez et al., 2013). *Marble burying*. Subjects were placed in mouse cages (NexGen, Allentown Inc., 194 mm x 181 mm x 398 mm) containing a 2 cm thick layer of corn husk bedding (Bed-O-Cobb) and sixteen marbles placed on top of the bedding. Subject’s activity was then videorecorded for 20 minutes and the number of marbles partially or fully buried at the end of the test were quantified. Marbles were considered to be fully buried if greater than 75% of the marble was submerged in the bedding. Marbles were considered partially buried if over 50% of the marbles was covered. Marbles were not considered to be buried if greater than 50% of the surface was exposed. Cages and marbles were cleaned with soap and water and dried between subjects. *Digging test*. Subjects were placed into a mouse cage (as above) containing a 2 cm thick layer of bedding for three minutes while videorecorded from above. The latency to start digging and the total duration of time spent digging were manually quantified from the recording. Cages were cleaned with soap and water and dried between subjects.

#### Forced swim

To determine whether iCre insertion altered active/passive coping behavior, we assessed each subject’s activity within the Forced Swim Test (Cryan et al., 2002; Cryan and Momereau, 2004). Subjects are placed into a cylindrical beaker (53 cm high, 23 cm diameter) filled with room temperature water to 17 cm depth for 5 minutes. The subject’s behavior was video-recorded from the side of the container, and the subject was continuously monitored to ensure they could stay afloat. At the end of the test, subjects were removed from the container, hand-dried, and then allowed to recover under a heating lamp in their home cage. The duration of time spent struggling/swimming and time spent floating were scored from the video for the last three minutes of the test using BORIS software (Friard and Gamba, 2016).

### Autoradiographic assessment of V1aR binding in *Avpr1a^Cre+^* knock-in mice

We measured V1aR binding in the lateral septum of *Avpr1a^Cre+^* knock-in mice (n=8 male, n=8 female) and their *Avpr1a^Cre-^* littermates as controls (n=8 male, n=8 female) to determine if iCre expression altered native expression of V1aR. In addition, tissue from wild-type mice (n=2 male, n=2 female) was processed for V1aR binding and was compared to matched brain sections from cre-reporter mice. Subjects were euthanized by carbon dioxide exposure and brains were immediately extracted and flash-frozen onto dry ice. Tissue was allowed to sit on dry ice for 20 min and was then stored at -80°C. Prior to sectioning, tissue was allowed to adjust to cryostat temperature (Leica CM3050 S, Leica Biosystems) for 30 min and was then coated in a thin layer of OCT compound and cut at 20 μm (coronal) sections. Sections were mounted directly onto slides at room temperature and allowed to dry prior to storage at -80°C.

Frozen slides with brain sections containing the LS were allowed to dry at room temperature for 30 min. Once dry, sections were carefully placed into metal slide racks and then fixed with 1% paraformaldehyde in a 0.1M phosphate buffer solution, followed by two 10-minute rinses with a 1X Trizma buffer solution (using Trizma crystals with preset pH). Sections were then incubated for 1 hr in tracer buffer (15% bacitracin and 50% bovine serum albumin in 1X Trizma buffer with MgCl) with iodinated linear vasopressin V1aR antagonist (Phenylacetyl 1, 0-Me-D-Tyr2, [125I-Arg6], (Perkin Elmer) to visualize V1aR binding, rinsed with 1X Trizma buffer with MgCl 3 times for 10 min each, and then briefly dipped in cold deionized water and dried with a hair dryer.

Dried slides were exposed to 35 cm x 43 cm Kodak film (Carestream BIOMAX 856 7232) for 72 hours. A carbon-14 standard (American Radiolabeled Chemicals) was placed in the center of each film for calibration. Films were developed and then imaged on an Epson Expression 12000XL photo scanner at 2400 dpi (8-bit grayscale).

Receptor binding was quantified and converted to disintegrations per minute (DPM/0.4 mg) by comparing it to the rate of decay of the carbon-14 standard using a NIH-Rodbard corrected normal curve generated by ImageJ (Shindelin et al. 2012). Image scale was set by measuring a scale box of known height (5 mm) and assigning the correct height in ImageJ. To control for background binding, density measurements were taken from cortical regions estimating nonspecific binding and this was subtracted from the specific binding for each measurement. The LS was identified on brain sections by comparing landmarks to relevant anatomical plates from Paxinos and Franklin (Fourth Edition, 2012).

### Assessment of cre expression in *Avpr1a^Cre+^* knock-in mice

*Avpr1a^Cre+^* mice were crossed with the Ai6 (Zs-Green; n=2/sex) or Ai14 (tdTomato; n=2/sex) cre-reporter lines (Madisen et al., 2009) to visualize iCre expression patterns driven by the *Avpr1a* gene throughout the lateral septum as well as other known V1aR-expressing brain regions. We also assessed the fidelity of *Avpr1a*-driven iCre expression by using *in situ* hybridization to determine the relationship between (a) *tdTomato* and *Avpr1a* mRNA expression in *Avpr1a^Cre+/tdTom^* subjects (n=2/sex) and (b) *iCre* mRNA and *Avpr1a* mRNA expression in *Avpr1a^Cre+^* subjects (n=2). As reporter expression reflects lifetime expression of iCre, we wished to determine the degree to which reporter expression in the LS reflects adult levels of iCre expression. To do so, we made bilateral stereotaxic injections of AAV-2/5-syn-FLEX-jGCaMP8m-WPRE (Addgene #162378-AAV5) into the LS of adult *Avpr1a^TdTom/Cre+^* subjects (n=2/sex) and processed brains for eGFP immunohistochemistry.

### *In situ* hybridization

We used non-radioactive *in situ* hybridization to determine the relationship between (a) *Avpr1a* and tdTomato mRNA expression in *Avpr1a^Cre+/tdTom^* subjects, (b) *Avpr1a* and *iCre* mRNA expression in *Avpr1a^Cre+^* subjects, (c) *Avpr1a* and oxytocin receptor (*Oxtr*), neurotensin (*Nts*), vesicular glutamate transporter 2 (*Slc17a6*), vesicular inhibitory amino acid transporter (*Slc32a1*), hypocretin receptor 1 (*hcrtr1)*, somatostatin (*Sst*), corticotropin-releasing hormone receptor 2 (*Crfr2*), and neuron-derived neurotrophic factor (*Ndnf*) mRNA transcripts in wild-type subjects.

Subjects were euthanized via carbon dioxide exposure, and their brains were immediately extracted and flash-frozen. Brains were then sectioned (20 μm thickness) on a cryostat (Leica CM3050 S) and mounted onto SuperFrost Plus slides (Fisher Scientific). Slides were fixed in 4% paraformaldehyde for 15 min at 4 °C, dehydrated in graded ethanol (50%, 70%, 100%), and air-dried. Hydrophobic barriers were drawn around tissue sections before probe application. *In situ* hybridization was performed using the RNAscope Multiplex Fluorescent v2 kit (Advanced Cell Diagnostics, Newark, CA, Cat. Nos. 323100, 323120) according to the manufacturer’s instructions with modifications optimized in our laboratory for fresh-frozen mouse brain tissue.

The following RNAscope probes were used: Mm-Oxtr-C3 (Cat. No. 412171-C3), iCre-C2 (Cat. No. 423321-C2), Mm-Avpr1a (Cat. No. 418061), Mm-Nts-C3 (Cat. No. 420441-C3), Mm-Slc17a6-C2 (Cat. No. 319171-C2), Mm-Slc32a1-C3 (Cat. No. 319191-C3), Mm-Hcrtr1-C2 (Cat. No. 466631-C2), Mm-Sst-C2 (Cat. No. 404631-C2), Mm-Crhr2-C3 (Cat. No. 413201-C3), and Mm-Ndnf-C3 (Cat. No. 447471-C3). Probes were hybridized for 2 h at 40 °C in the HybEZ II oven (Advanced Cell Diagnostics, Cat. No. 321720). Following probe hybridization, signal amplification steps were carried out sequentially using Amp 1-3 reagents and horseradish peroxidase (HRP) channel development according to the RNAscope Multiplex Fluorescent v2 workflow. TSA fluorophores (Advanced Cell Diagnostics; TSA Vivid 520, 570, and 650, Cat. Nos. 323271, 323272, and 323273, respectively) and Akoya Biosciences Opal fluorophores (Opal 520, 570, and 690, Cat. Nos. FP1487001KT, FP1488001KT, and FP1497001KT, respectively) were applied at dilutions optimized for probe expression levels using RNAscope Multiplex TSA buffer (Advanced Cell Diagnostics; Cat. No. 322810). Between amplification steps, slides were washed twice in the RNAscope wash buffer (Advanced Cell Diagnostics; Cat. No. 310091) as recommended. After the final amplification and HRP-blocking steps, slides were counterstained with DAPI and coverslipped with ProLong Gold antifade mounting medium (Thermo Fisher Scientific, Cat. No. P36930). Sections were stored at 4 °C until imaging.

### Immunohistochemistry

Subjects were euthanized using pentobarbital-based euthanasia solution (150mg/kg) and perfused with phosphate-buffered saline (PBS) followed by 4% paraformaldehyde (PFA) solution. Brains were extracted and immediately drop-fixed in 4% PFA solution for at least 12 hours and then transferred to a 30% sucrose solution for at least 48 hours. The brain was then sectioned at 30μm and stored in an antifreeze cryoprotectant until immunohistochemical processing. The tissue was rinsed in PBS, then incubated at room temperature overnight in a primary antibody solution (0.04% Triton-X) for anterograde (synaptophysin) tracing (1:1000 rabbit anti-RFP, abcam ab124754; 1:5000 chicken anti-GFP, abcam ab13970) and for monosynaptic retrograde (rabies) tracing (1:750 chicken anti-GFP, abcam ab13970). Tissue from AAV-eGFP injections in tdTomato mice were also incubated with 1:5000 chicken anti-GFP primary antibody (abcam ab13970). Sections were rinsed in PBS and incubated at room temperature for 2 hours in a fluorescent secondary antibody solution containing 0.04% Triton-X for anterograde tracing experiments (1:600 goat anti-rabbit, Alexa Fluor 594, abcam ab150080; 1:600 goat anti-chicken, Alexa Fluor 488, abcam ab150169) and for retrograde tracing experiments (1:500 goat anti-chicken, Alexa Fluor 488, abcam ab150169). Sections were then rinsed in PBS and mounted onto SuperFrost Plus slides (Fisher Scientific) and cover-slipped with Prolong Gold with DAPI (Life Technologies, Carlsbad, CA, USA). To differentiate hippocampal CA2 cells from hippocampal CA1/CA3 cells, we utilized a combination of anti-PCP4 (1:400, rabbit anti-PCP4, Sigma Aldrich, HPA005792) and anti-RGS14 (1:250, mouse anti-RGS14, antibodies inc., AB_2179931) primary antibodies to target the dorsal and ventral CA2 respectively, along with corresponding secondaries (1:500 goat anti-rabbit, Alexa Fluor 488, ab150077; 1:500 goat anti-mouse, Alexa Fluor 488, ab150113), as described above.

### Identifying LS*^Avpr1a^* cell connections

#### Anterograde tracing

We identified the neural targets of LS*^Avpr1a^* cells by placing unilateral stereotaxic injections of an anterograde adeno-associated virus (AAV) tracer (hSyn-Flex-mGFP-2A-Synaptophysin-mRuby; Addgene #71760) into the LS of *Avpr1a^Cre+^* (N9 generation) males (n=1/sex) and females (n=1/sex) as well as *Avpr1a^Cre-^* subjects (n=2/sex) as controls. This vector expresses, in a cre-dependent manner, membrane-bound GFP in axons and mRuby in putative presynaptic terminals (Beier et al., 2015). The side injected was randomized between animals to reduce any possible hemisphere bias. Six weeks later, subjects were sacrificed and processed for immunohistochemical analysis for LS*^Avpr1a^* output fiber and presynaptic terminal location.

#### Retrograde tracing

We utilized a modified rabies virus strategy to investigate the monosynaptic inputs to LS*^Avpr1a^* cells (Kim et al., 2016, Lavin et al., 2019) of male and female *Avpr1a^Cre+^* mice (N9 generation; n=1/sex) as well as littermate *Avpr1a^Cre-^* controls (n=1/sex). This approach uses G-deleted rabies virus (RV, SADB19; RVdG-mCherry; Kavli Institute for Systems Neuroscience, Trondheim, Norway), in which the G gene, which encodes the glycoprotein needed for cell entry, is replaced with mCherry fluorophore and then pseudotyped with an avian sarcoma leukosis virus glycoprotein (EnVA), which prevents viral infection of mammalian cells. Both the G protein and the entry protein for EnVA (TVA) were introduced to LS*^Avpr1a^* cells via unilateral (alternate sides) stereotaxic injection of two AAV helper viruses: AAV2/1-syn-FLEX-splitTVA-EGFP-tTA (tTA; Addgene #100798), which expresses TVA, eGFP, and tetracycline trans-activator (tTA) in a Cre-dependent manner, and AAV2/1-TREtight-mTagBFP2-B19G (G; Addgene #100799), which expresses G and blue fluorophore under control of a tetracycline response element (Liu et al., 2017). The tTA virus Cre-dependently expresses TVA, which allows the EnVA-pseudotyped RV to infect LS*^Avpr1a^* cells. tTA expression also drives the expression of the co-expressed G, allowing RV to produce the envelope protein G that allows transsynaptic transport. Ten days after injection of the helper viruses (250 nl; 1:1 mixture of tTA and G; G virus diluted 1:20 in Dulbecco’s phosphate-buffered saline; Fisher, 14–190-250) into the LS, we injected RV (200 nl) into the same location. Seven days later, subjects were sacrificed and processed for immunohistochemical or *in situ* hybridization analysis.

### Imaging and analysis

#### Imaging

Images of RNAscope tissue, synaptophysin puncta counts, and starter cell location were taken at 10x, 20x, and 40x using Zeiss LSM 700 laser scanning confocal (Carl Zeiss Microimaging, Göttingen, Germany) with Zen Microscopy Software (Zen 2012 SP1 black edition V8.1). Tissue from reporter mice as well immunohistochemical material were imaged at 5x on a Zeiss Axio Imager.M2 microscope (Carl Zeiss Microimaging, Göttingen, Germany), which transferred fluorescent images (FITC contrast reflector) to image analysis software (Stereo Investigator, MicroBrightField, RRID:SCR_002526). Bilateral imaging domains were placed with reference to anatomical landmarks such as ventricles. Additionally, tiled whole section images were taken at 10x magnification using a Zeiss Inverted Axiovert 100M (Carl Zeiss Microimaging, Göttingen, Germany) and Micro-Manager open source software (Edelstein et al., 2014). Tiling was performed using the FIJI Stitching plugin (Preibish et al., 2009; Shindelin et al., 2012).

#### Analysis

##### RNAscope

All RNAscope-labeled tissue sections were analyzed using QuPath software (version 0.4.4) (Bankhead et al., 2017). For each RNAscope experiment, we selected one section per animal that contained the posterior dorsal (LSd) and intermediate regions of LS (LSi) and showed robust signals for the target probes. Cell segmentation was performed on defined LSd and LSi regions of interest using the DAPI channel to identify nuclear and cellular boundaries. Within QuPath, we used the Cell Detection function to generate cell segmentations. Segmentation parameters (i.e. background radius, threshold, and cell expansion) were manually optimized for each RNAscope probe set to achieve accurate delineation of individual cells across experiments. For mRNA puncta detection, we used the Subcellular Detection tool within QuPath. This allowed us to quantify the number of fluorescent puncta corresponding to each mRNA probe within individual cell boundaries. Detection thresholds were adjusted for each probe channel to minimize background and maximize true signal. Following detection, measurement data for each cell (including cell ID, area, and puncta counts for each channel) were exported from QuPath to Microsoft Excel for further analysis. The exported data were combined across both left and right sides of each section and across all animals for downstream quantification. Cells were considered positive for a transcript if they contained more than one punctum for that probe. This puncta cutoff differed across probes such that low expression targets (V1aR, Hcrt) required a lower threshold of 2+ puncta, whereas higher expressing targets had higher thresholds (CRFR2, 3+ puncta; SST, NDNF, NTS, GFP, 4+). Imaging thresholds were optimized separately for each probe to account for differences in hybridization efficiency and signal intensity.

##### Anterograde tracing (synaptophysin)

Tissue was imaged using multichannel immunofluorescence microscopy to identify virally infected LS cells and their GFP-labeled fibers and synaptophysin–mRuby–labeled putative synaptic terminals. Fiber locations were identified by reference to anatomical plates in the Paxinos and Franklin mouse brain atlas (Fourth Edition, 2012) and fiber density was scored semiquantitatively: +, very few fibers; ++, few dispersed fibers; +++, part of region densely covered by fibers; ++++, most of region densely covered by fibers (Rigney et al, 2023). Brain regions expressing both GFP-labeled fibers and mRuby-labeled putative synaptic terminals were considered direct outputs of V1aR LS cells. Regions with high expression of fibers were imaged at 40x and puncta overlapping with fibers were scored semiquantitatively: +, very few to no puncta; ++, few dispersed puncta; +++ some puncta; ++++ dense puncta across region.

##### Retrograde tracing (rabies)

The tissue was imaged using multichannel immunofluorescence microscopy to identify virally infected cells by brain region using the Paxinos and Franklin mouse brain atlas (Fourth Edition, 2012). LS cells from which the RV has spread (“starter cells”) will be identified by coexpression of GFP from the tTA virus and mCherry from RV and quantified. Then, input cells, identified by expression of exclusively mCherry from the RV, were identified by brain region, imaged at 10x and 20x, and cells on every other section were counted. Input cells by region were semiquantitatively ranked: + 1-2 cells; ++, 3-4 cells; +++, 5-6 cells; ++++, 7+ cells.

#### Statistical analysis

All data were analyzed and graphed in R (R Studio v4.3.2 2023-10-31; Windows; R Core Team, 2024). Packages used for data analysis were tidyverse, ggplot2, rstatix, emmeans, ggpubr, car, afex, and multcomp. If data violated the assumption of homogeneity of variance, nonparametric tests were used. All health assessments, sensorimotor assessments, and behavioral assessments were analyzed using a 2-way ANOVA with subject sex and genotype (Cre+ or Cre-) as between factors. Post-hoc Tukey HSD tests were used to test for differences across genotypes and sex, with a p-value adjustment for multiple comparisons. Effect sizes were calculated using partial eta-squared (ηp^2^) for all significant ANOVA effects. Autoradiographic binding density was analyzed using a linear mixed effects model.

## Results

### Behavioral and anatomical validation of *Avpr1a^Cre+^* mice

#### Behavioral assessment

*Avpr1a^Cre+^* mice were compared to their iCre-littermates on a variety of basic health assessments: vibrasse condition, body weight, and body length. No significant difference was found between genotypes in any health assessment in either sex (Table 1). iCre+ and iCre-littermates were also compared for sensorimotor function: visual placing, contact placing, limb clasping, auditory function, grip strength, olfaction, and motor function (rotarod). No significant difference was found between genotypes of either sex in any of these assessments (Table 1). Finally, iCre+ and iCre-littermates underwent a series of behavioral assessments examining activity levels, anxiety- and depression-like behaviors, and reward processing: elevated plus maze, open field, marble burying, bedding digging, forced swim test, and the sucrose preference test. None of the behavioral assessments revealed any significant difference between genotypes (Table 1).

#### Anatomical assessment

To assess the accuracy of Cre expression in the *Avpr1a^Cre^* line (homozygous subjects), we performed RNAscope *in situ* hybridization to compare *iCre* and *Avpr1a* mRNA expression within LS. Imaging and analysis were done on the posterior LS, specifically the dorsal (LSd) and intermediate (LSi) subdivisions, which were selected for their strong and consistent V1aR signal. Within these regions, Cre and V1aR mRNA showed substantial overlap, with almost 100% of cells co-expressing both transcripts, indicating that the knock-in construct effectively drives Cre expression in V1aR+ cells (Figure 2A). To ensure that insertion of the P2A-iCre cassette did not alter receptor functionality, we compared the autoradiographic assessment of V1aR binding in *Avpr1a^Cre+^* mice and their Cre-littermates. V1aR binding density within the LS and other highly expressed brain regions did not differ by genotype (F_1,29.63_ = 1.75, p = 0.20), sex (F_1,31.96_ = 1.01, p = 0.32) or the interaction between genotype and sex (F_1,29.63_ = 2.60, p = 0.12) (Figure 2B).

**Figure 2.**
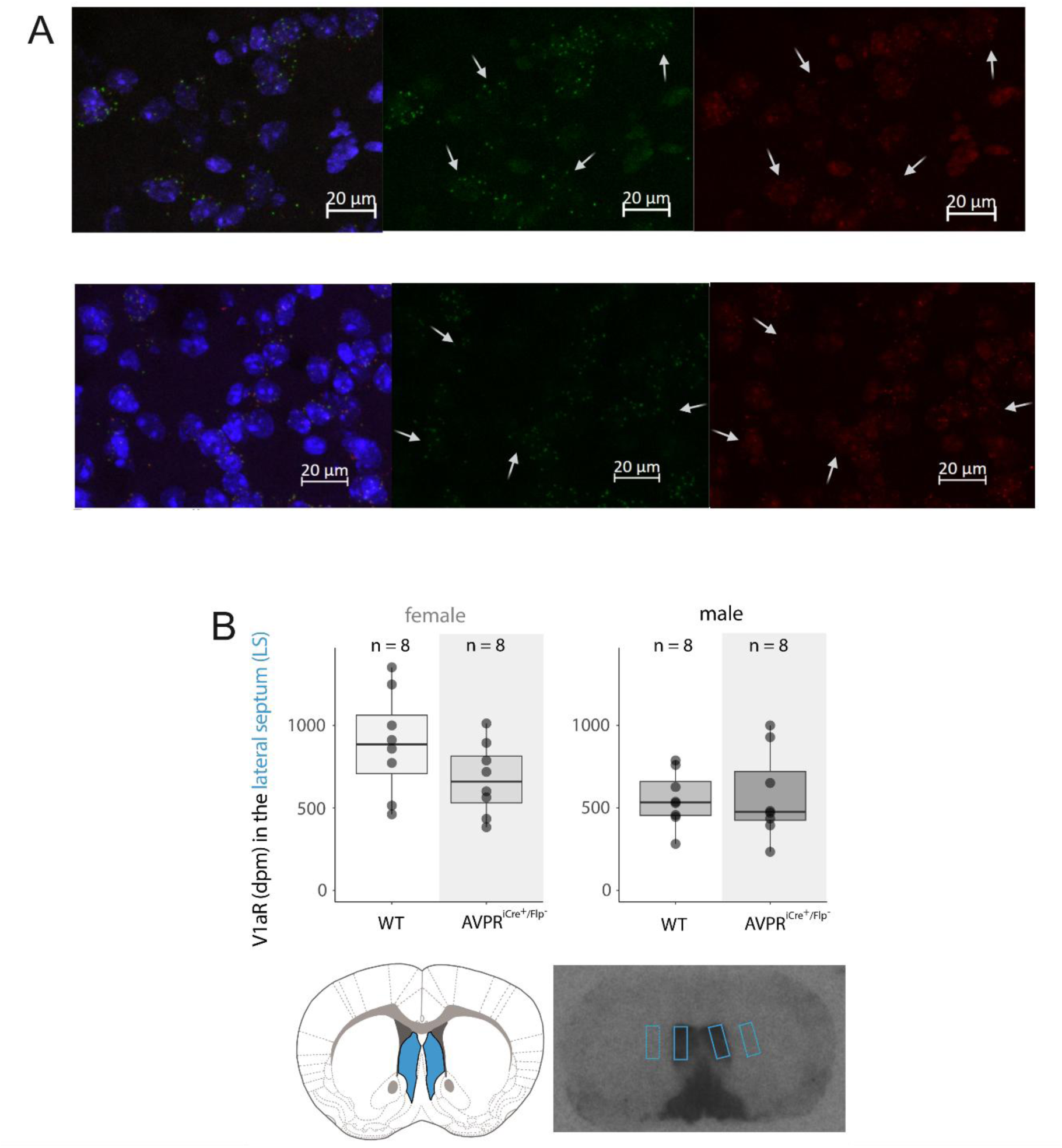
Confirmation of V1aR expression and colocalization with iCre in the *Avpr1a^Cre+^* strain **A.** *In situ* hybridization confirmation of V1aR and iCre colocalization in the *Avpr1a^Cre+^* strain. Blue is DAPI, green indicates V1aR mRNA, and red indicates iCre mRNA, arrows indicate cells which are V1aR+ and iCre+. **B.** Autoradiograph of V1aR binding density comparing *Avpr1a^Cre+^* and wildtype littermates. No difference was found in binding density in males or females.

To verify the fidelity of *Avpr1a*-driven iCre expression, we crossed *Avpr1a^Cre+^* mice with reporter lines (Ai6, ZsGreen; Ai14, tdTomato). Reporter expression in *Avpr1a^Cre+/ZsGreen^* and *Avpr1a^Cre+/tdTom^* mice showed strong labeling in brain regions previously identified as sites of high V1aR expression, including the lateral septum (LS), diagonal band of Broca (DBB), and lateral hypothalamus (LH) (Figure 3). The distribution of fluorescence was similar to the pattern of V1aR mRNA expression reported in the Allen Brain Atlas, showing that the Cre knock-in closely matches natural *Avpr1a* expression (*Experiment #74641316*, Allen Institute for Brain Science; Lein et al., 2007). Reporter patterns were consistent across sexes and between reporter lines (Figure 3C-L).

**Figure 3.**
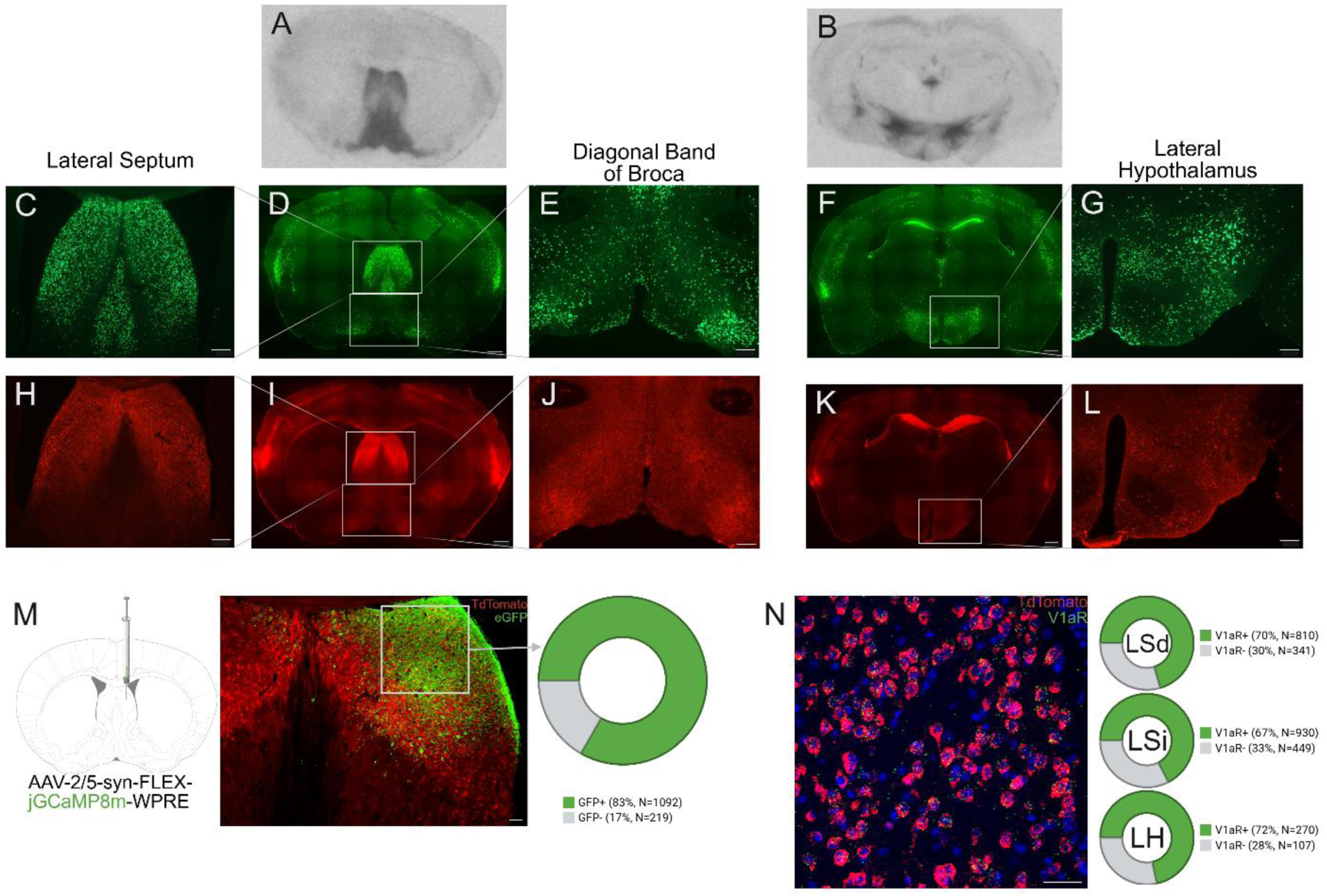
Validation of *Avpr1a^Cre+^* strain through crosses with reporter strains expressing cre-dependent ZsGreen and TdTomato. **A-B.** Autoradiograph of V1aR binding density in wildtype male mouse. Sections are approximately +0.50mm (A) and - 1.82mm (B) from bregma. **C-G.** Examples of tag expression from *Avpr1a^Cre+/ZsGreen^* male mouse. Known regions of high V1aR expression were selected and imaged at 5x (C, lateral septum; E, diagonal band of broca; G, lateral hypothalamus; scale bar = 100μm) while sections containing these entire regions were imaged at 10x and tiled (D and F; scale bar = 500μm). Sections are approximately +0.50mm (C-E) and -1.83mm (F-G) from bregma. **H-L.** Examples of *Avpr1a^Cre+/tdTom^* male mouse V1aR expression. Known regions of high V1aR expression were selected and imaged at 5x (H, lateral septum; J, diagonal band of broca; L, lateral hypothalamus; Scale bar = 100μm) while entire sections containing these regions were imaged at 10x and tiled (I and K; Scale bar = 500μm). Sections are approximately +0.50mm (H-J) and -1.83mm (K-L) from bregma. **M.** Representative image showing tdTomato reporter fluorescence (red) and AAV2/5-syn-FLEX-jGCaMP8m-driven eGFP expression (green) in the lateral septum of adult *Avpr1a^Cre+/tdTom^* mice, imaged at 5x (scale bar = 50μm). eGFP immunolabeling marks cells with active Cre recombination following viral injection. Donut charts summarize the percentage of tdTomato^+^ cells co-labeled with eGFP across lateral septum. **N.** Representative *in situ* hybridization image from *Avpr1a^tdTom/cre+^* mice showing tdTomato mRNA (red) and V1aR mRNA (green) expression within the lateral septum, imaged at 20x (scale bar = 50μm). Donut charts show the percentage of tdtomato^+^ cells containing V1aR mRNA puncta across dorsal (LSd), intermediate (LSi), and lateral hypothalamus (LH).

To confirm that reporter expression in *Avpr1a^Cre^* mice reflects adult *Avpr1a* expression, we injected AAV-2/5-syn-FLEX-jGCaMP8m bilaterally into the LS of adult *Avpr1a^Cre+/tdTom^* mice to compare reporter labeling with Cre-dependent viral expression. Expression of jGCaMP8m, visualized by eGFP immunohistochemistry, showed a strong overlap with tdTomato reporter expression from developmental Cre activity (Figure 3M), with 83% of tdTomato⁺ cells co-labeled with eGFP (1,311 total cells analyzed). *In situ* hybridization (ISH) further confirmed that tdTomato expression in *Avpr1a^Cre+/tdTom^* mice was largely colocalized with *Avpr1a* mRNA within the lateral septum (LS) (Figure 3N). Quantitatively, 70% of tdTomato⁺ cells in LSd (1,151 total cells), 67% in LSi (1,379 total cells), and 72% in LH (134 total cells) contained V1aR mRNA puncta, indicating that reporter expression reflects endogenous V1aR transcription.

### Anterograde outputs from LS V1aR+ cells

To identify outputs of LS V1aR cells we unilaterally injected Cre-dependent hSyn-Flex-mGFP-2A-Synaptophysin-mRuby into the LS of male and female *Avpr1a^Cre+^* mice (Figure 4A). In Cre-negative controls we saw no viral expression. In the Cre-positive male and female, viral spread, as identified by GFP-labeled cell bodies, was limited to the LS, primarily the LSi (Figure 4B).

**Figure 4.**
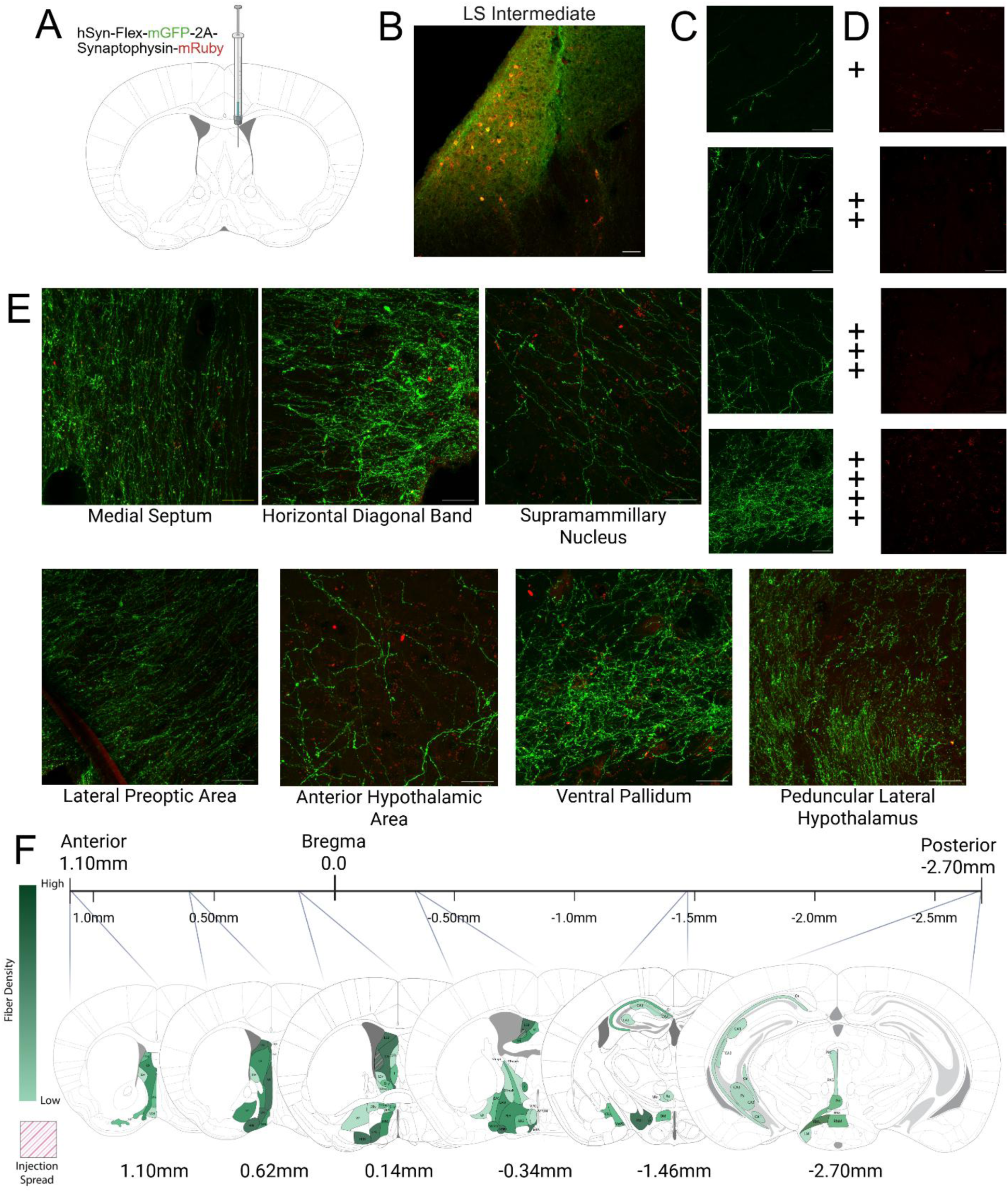
Anterograde tracing of LS V1aR cells. **A.** Representative image of unilateral injection of Synaptophysin AAV into the LS (10x, Scale bar = 50μm). **B.** Example of injection spread in the LS. **C.** Example images of fiber densities for each semiqualitative ranking: +, very few fibers; ++, few dispersed fibers; +++, part of region densely covered by fibers; ++++, most of region densely covered by fibers (40x, Scale bars = 50μm). **D.** Example images of puncta densities for each semiqualitiative ranking: +, very few to no puncta; ++, few dispersed puncta; +++ some puncta; ++++ dense puncta across region (40x, Scale bars = 50μm). **E.** Examples of fibers and puncta in regions of interest in one female brain: medial septum, horizontal diagonal band, lateral preoptic area, anterior hypothalamic area, ventral pallidum, penducular lateral hypothalamus, and supramammillary nucleus (40x, Scale bars = 50μm). **F**. Representative diagram of fiber density and injection spread throughout one female brain.

#### Telencephalon

There were small amounts of fibers seen throughout the dorsal and ventral hippocampal CA1, CA2, and CA3 regions. LS V1aR cells had strong projections to, as evidenced by high density of puncta, the ventral pallidum (VP) and the diagonal band of broca (DBB; Figure 4E). In the VP, the strength of innervation was higher anterior than posterior while the DBB had strong inputs throughout. The shell of the nucleus accumbens showed the most anterior innervation, though fiber density was minimal. As expected, the bed nucleus of the stria terminals (BNST) and the medial amygdala (MeA), both of which are major AVP inputs to the lateral septum, displayed projections from LS V1aR cells throughout each structure, with fibers and puncta denser in the MeA than the BNST. Most projections were ipsilateral, but some less dense contralateral projections were found primarily in the LS opposite the injection site and other septal structures such as the medial septum, septohippocampal nucleus, and the lambdoid septal zone (Figure 4E). Little to no fibers were seen in the cerebral cortex.

#### Diencephalon

Thalamic regions such as the reticular thalamus, paraventricular nucleus (PVN), and intermediodorsal nucleus received sparse occasional projections with relatively dense puncta while the nucleus of reuniens had light fibers throughout much of its structure. The hypothalamus was one of the strongest outputs for LS V1aR cells. Hypothalamic regions which exhibited dense fibers and puncta include the ventral lateral hypothalamus, peduncular lateral hypothalamus, magnocellular nucleus of the lateral hypothalamus, anterior hypothalamic area, medial preoptic area, medial and lateral preoptic nucleus, and the supramammillary nucleus (Figure 4E). Contralateral projections were found primarily in the supramammillary nucleus.Minimal fibers were found in the lateral and medial habenula.

#### Mesencephalon

Caudal LS V1aR projections were limited in the mesencephalon. Fibers were primarily found ipsilaterally in small amounts in the medial periaqueductal gray.

### Monosynaptic inputs to LS V1aR+ Cells

To identify inputs to LS V1aR cells we utilized a G-deleted rabies virus (RVdG-mCherry) injected unilaterally following cre-dependent helper viruses (AAV2/1-syn-FLEX-splitTVA-EGFP-tTA and AAV2/1-TREtight-mTagBFP2-B19G) in *Avpr1a^Cre+^* and Cre-negative male and female mice (Figure 5A-B). Cre-negative mice showed no RV expression. The Cre+ male and female showed starter cells, as identified by coexpression of GFP and RFP, exclusively in the LS, across the LSd, LSi, and LSv. (Figure 5D).

**Figure 5.**
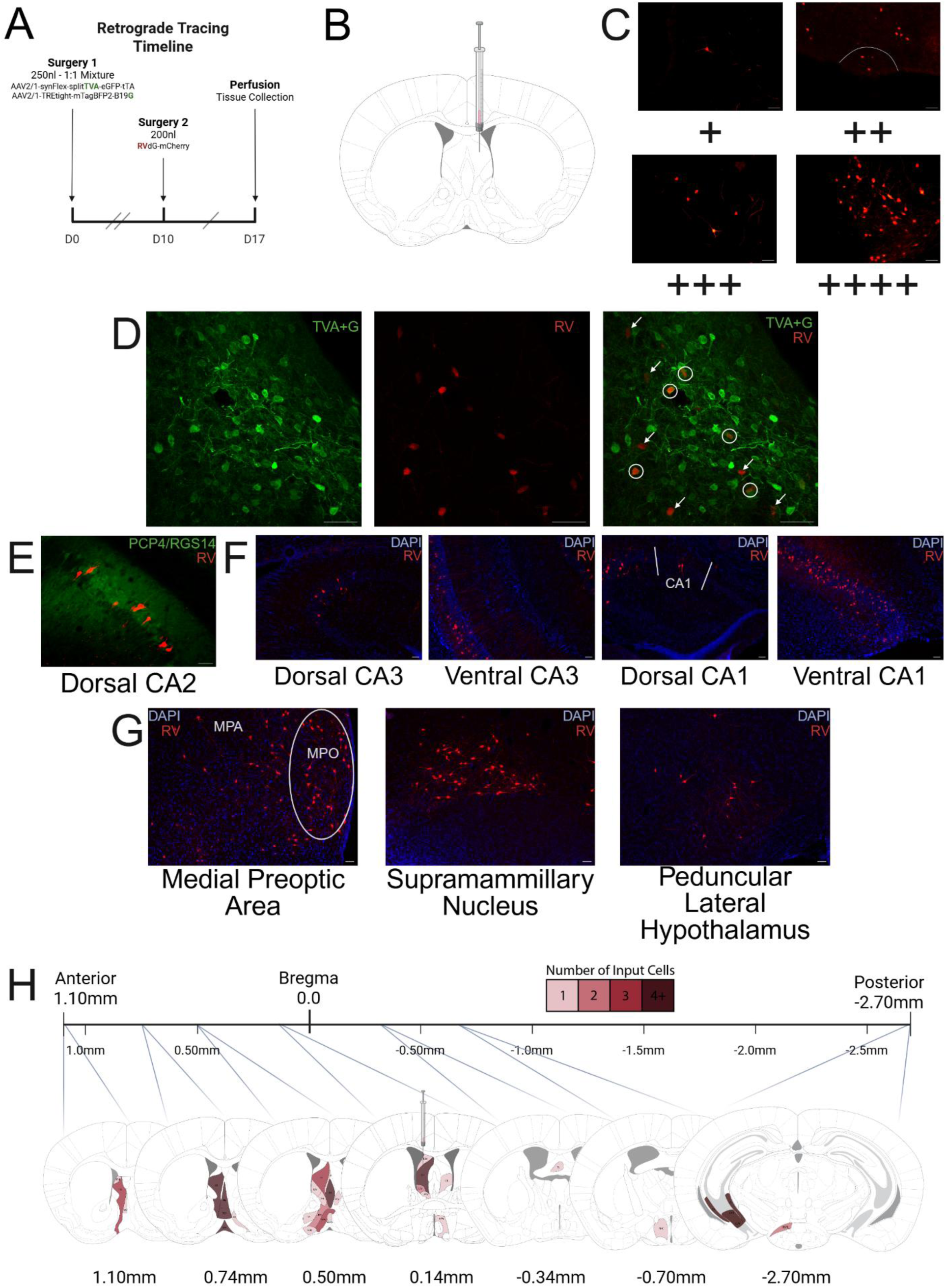
Retrograde tracing of LS V1aR cells. **A.** Timeline for injections of TVA, G, and RV viruses. **B.** Representative image of unilateral injection of TVA, G, and RV viruses into the LS.. **C.** Example images of input cell densities for each semiqualitative ranking: + 1-2 cells; ++, 3-4 cells; +++, 5-6 cells; ++++, 7+ cells. (20x, Scale bars = 50μm). **D.** Example of starter cells in the LS. Green indicates TVA+G virus and red indicates RV. Cells expressing both TVA+G and RV are considered starter cells while cells expressing just RV are input cells. (20x, Scale bars = 100μm) **E.** Example of input cells in dorsal CA2. Green indicates PCP4/RGS14 and red indicates RV. **F.** Examples of input cells in Dorsal/Ventral CA3 and Dorsal/Ventral CA1. Blue indicates DAPI and red indicates RV. (20x, Scale bars = 50μm). **G.** Examples of inputs from notable regions: medial preoptic area, supramammillary nucleus, and peduncular lateral hypothalamus (20x, Scale bars = 50μm). **H.** Representative diagram of input cell density throughout one female brain.

#### Telencephalon

Input cells were found all throughout the VP and the DBB, particularly the VDB. The BNST and MeA also had many input cells. This is not surprising as these regions are the primary sources of AVP to the LS and previous studies have shown strong output from these regions to the LS (Caffé et al., 1987; De Vries and Buijs, 1983; Rigney et al., 2023). Many input cells were also found in the MS and the LS opposite to the injection. Dorsal and ventral hippocampal CA1, CA2, and CA3 also contained many input cells, the most of any region. CA3 had the strongest output to the LSV V1aR cells, followed by CA2, then CA1 (Figure 5E-F). Most regions which had strong input ipsilaterally had lesser input cells contralaterally. No input cells were found in the cortex.

#### Diencephalon

Minimal input cells were found in the medial and lateral habenula. Hypothalamic regions overall contained large numbers of input cells. Some of the highest input cell numbers come from regions such as medial preoptic area, lateral preoptic area, medial preoptic nucleus, peduncular lateral hypothalamus and the supramammillary nucleus (Figure 5G). Most of these regions also receive dense projections from the lateral septum (see above). Most thalamic areas such as the reticular thalamus contained a few input cells. However, the paraventricular nucleus of the thalamus contained many input cells across its substructure. As in the telencephalon, most regions with dense input cells ipsilaterally have contralateral input cells to a lesser degree.

#### Mesencephalon

Overall, few input cells were found in mesencephalic areas. Small amounts were found in the ventral tegmental area and the medial periaqueductual gray. All of these input cells were ipsilateral to the injection site.

### Characterizing V1aR cells in the LS

All V1aR+ cells in LS co-expressed *Vgat*, a GABAergic cell marker (Supplementary Figure 1), but not *Vglut2*, a glutamatergic marker, indicating that essentially all V1aR cells in LS are inhibitory. In comparing V1aR expression to that of Oxtr throughout the LS, we found increasing expression of V1aR in both the posterior and dorsal axes of the LS. In contrast, Oxtr expression seems largely consistent across the anterior-posterior axis, and towards more ventral regions of the LS. The LSi, especially the posterior region, had substantial V1aR/Oxtr coexpression, with no obvious sex differences (Figure 6B-C).

**Figure 6.**
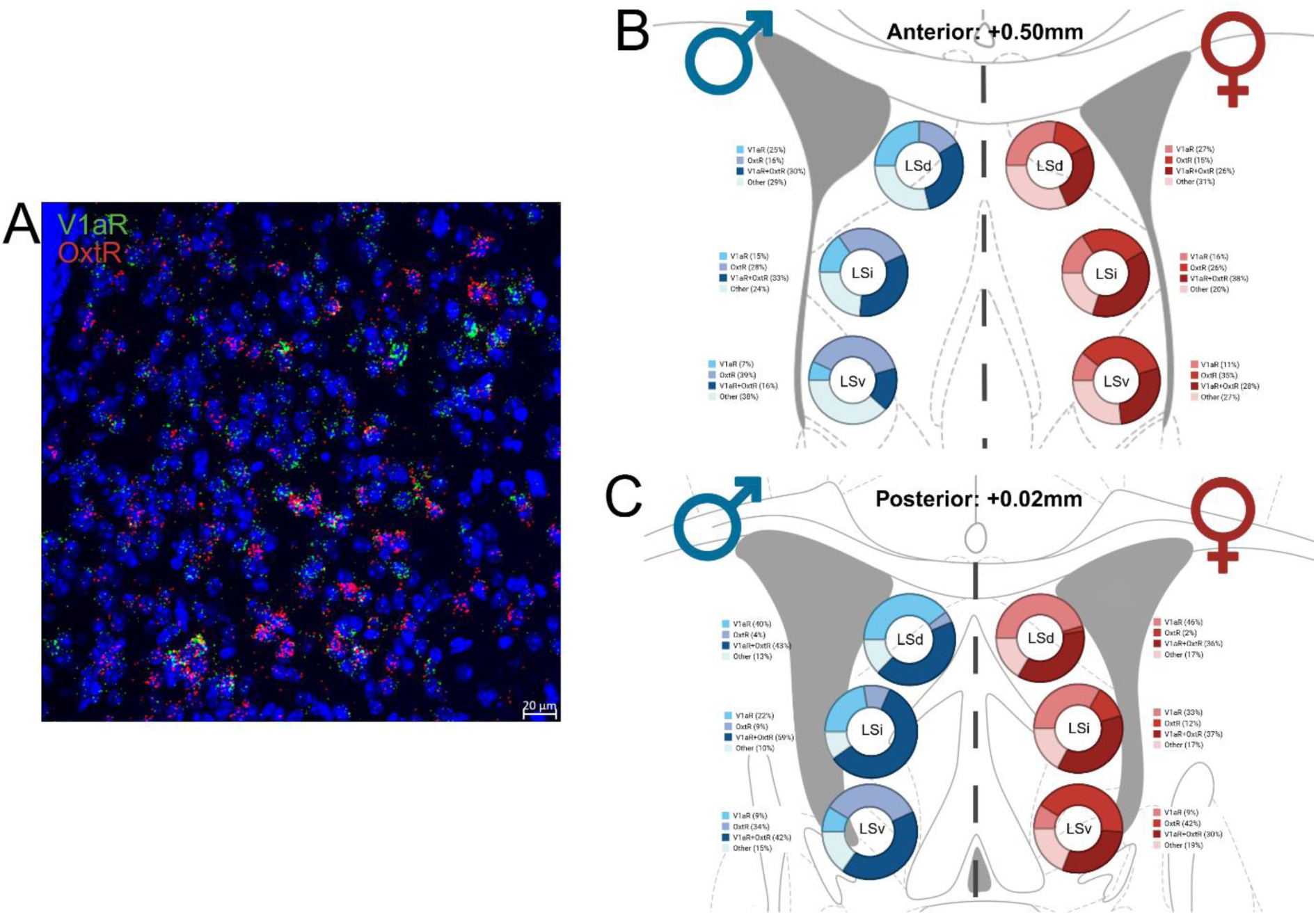
*In situ* hybridization Oxtr and V1aR colocalization in the LS. **A.** Example image of the colocalization of Oxtr mRNA (red) and V1aR mRNA (green; blue indicates DAPI; 20x). **B-C.** Percentages of cells in male and female brains expressing Oxtr only, V1aR only, both, or neither in the anterior and posterior LSd, LSi, and LSv.

We noted that *Crfr2* expressed at high levels in LSi and was strongly colocalized with V1aR in that subregion, with approximately 2/3 of *Crfr2+* cells expressing V1aR puncta and half of V1aR+ cells expressing *Crfr2*. In the LSd, there were few *Crfr2+* cells and so only 5% of V1aR+ cells in this region expressed *Crfr2*. Although *Hcrtr1* was expressed in only a small number of LSd and LSi cells (∼1%), these cells were highly co-expressed with V1aR puncta (70%). *Sst* expression was higher in LSd cells (18%) than LSi (8%) cells, and showed much higher co-expression with V1aR in the LSd (87%) than in the LSi (57%). V1aR+ cells also were much more likely to be positive for somatostatin in the LSd (26%) than in the LSi (9%). *Nts* expression was more prevalent in LSi (7%) than LSd (5%), but exhibited higher coexpression with V1aR in the LSd (71%) than in the LSi (53%), while V1aR expression of *Nts* was only 5% and 6% in LSd and LSi respectively. *Ndnf* coexpresses prominently with V1aR cells of both the LSd (80%) and LSi (83%), but *Ndnf* expression is much higher in the LSd, with 90% of V1aR+ cells expressing *Ndnf* in the LSd compared to 21% in the LSi. Percentages of total cells containing V1aR, percentage of total containing above-mentioned markers (*Crfr2*, *Hcrtr*, *Ndnf*, *Nts*, or *Sst*), and percentage of total cells containing both markers or neither marker are represented in Figure 7.

**Figure 7.**
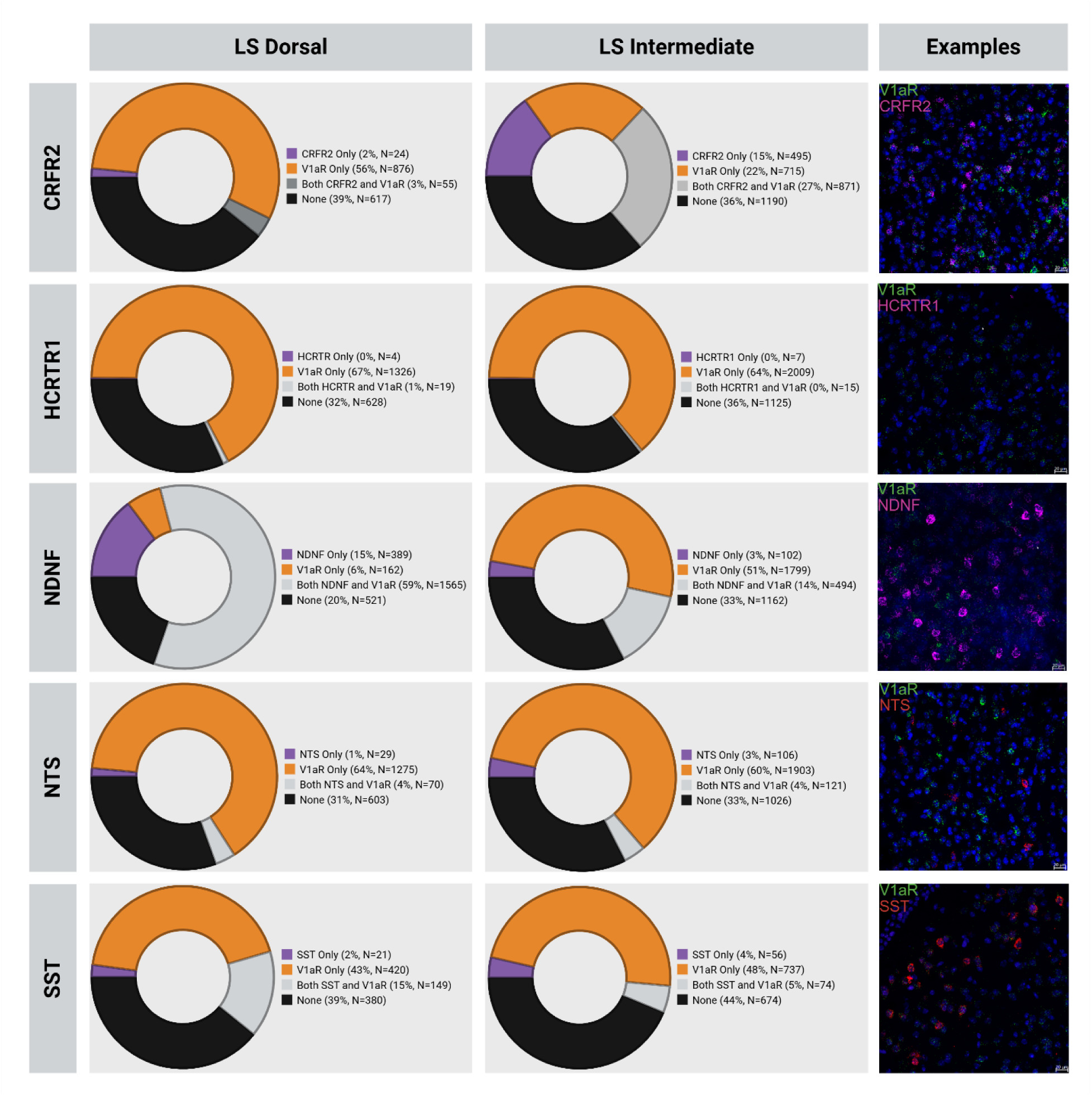
*In situ* colocalization percentages of V1aR with multiple genes (CRFR2, HCRTR1, NDNF, NTS, SST) in the dorsal and intermediate LS. Representative images in the LS show V1aR (green) colocalized with CRFR2 (magenta), HCRTR1 (magenta), NDNF (magenta), NTS (red), and SST (red; blue indicates DAPI; 20x).

## Discussion

In this study we demonstrated that our V1aR Cre knock-in mouse line was phenotypically similar to wild-type mice and that brain Cre expression pattern matched known V1aR binding patterns as well as colocalizing with V1aR mRNA. Through a variety of health, physiological, and behavioral assessments we confirmed that the strain does not differ significantly from wildtype littermates. Using autoradiography, we confirmed that the Cre insertion in the *Avpr1a* gene does not alter V1aR ligand binding. We confirmed, through both reporter strain crosses with *Avpr1a^Cre+^* and Cre-dependent AAV injection into LS of *Avpr1a^Cre+/tdTom^* mice, that Cre expression mirrors endogenous V1aR expression. ISH also confirmed that tdTomato expression in *Avpr1a^Cre+/tdTom^* mice was largely colocalized with *Avpr1a* mRNA within the LS, indicating that reporter expression reflects endogenous *Avpr1a* transcription. This finding indicates that Cre recombinase remains active in adulthood and continues to be expressed in the same LS neuronal populations identified in the reporter line crosses. Together, these results demonstrate that the *Avpr1a^Cre^* line reliably captures both developmental and adult *Avpr1a* expression patterns, making it a robust tool for functional studies of V1aR-expressing neurons. However, we did notice that there was greater expression of tdTomato/ZsGreen signal in reporter mice in both canonical V1aR-expressing areas as well as other brain regions, such as the hippocampus, that do not show autoradiographic V1aR expression in adulthood. This is not surprising as tag expression in Cre-reporter mice marks all post-conceptional Cre-expressing cells, including cells in adulthood as well as those during development (Dymecki and Kim, 2007). Indeed, V1aR expression has been observed in cortex and hippocampal CA1 in neonatal, but not adult mice (Hammock & Levitt, 2012). The development and validation of this *Avpr1a^Cre+^* mouse line makes possible genetic access to a neuropeptide receptor system important for physiology and social, emotional, and mnemonic processes (Rigney et al., 2022).

We used Cre-dependent viral tracing approaches to examine the connectivity of LS V1aR neurons. Anterograde AAVs expressing membrane-bound GFP and synaptophysin (RFP) reporters revealed LS V1aT efferent projections and their potential terminal synapses (Bier et al., 2015; Rigney et al., 2023; Zingg et al., 2017). Complementary monosynaptic rabies-based retrograde tracing identified direct presynaptic partners of LS V1aR cells (Wickersham et al., 2007; Ottenheimer et al., 2018; Rigney et al., 2023). Both the anterograde and retrograde tracing are consistent with previous work in rats and mice (Caffé et al., 1987; DeVries and Bujis, 1983; Rood et al., 2013). Many regions which received strong projection from LS V1aR cells also had large quantities of input cells to LS V1aR cells (Figure 4E-F; Figure 5G-H). Several of these regions, such as VP, DBB, LH, and SuM, also contain V1aR cells themselves and are known to receive input from extended amygdala AVP cells (Figure 3A-B; Hammock and Levitt, 2012, Hernandez et al., 2016, Rigney et al., 2023). This suggests there is an integrated AVP circuit tightly binding together V1aR cell activity throughout the brain. Future work should target outputs of V1aR LS cells to investigate this circuit further. Specific targeting of the subregions of the LS (LSd, LSi, LSv) will provide more insight into this circuitry.

The primary input to LS V1aR cells are the hippocampal CA1, CA2, and CA3 regions. The hippocampus is known to have a role in social behavior, CA3 in particular has been implicated in social memory in mice (Chiang et al., 2018). CA2 is of particular interest as it is one of the few regions in the brain which contains the vasopressin 1b receptor (V1bR; Young et al., 2007). AVP action on the V1bR present on CA2 terminals within the LS has been shown to influence social aggression in mice (Leroy et al., 2019). We have confirmed that the CA2 contains input cells to the LS. Whether these cells contain V1bR remains uncertain. If the CA2 input cells are the same V1bR cells which project to the LS, then AVP release in the LS may act on both V1bR on presynaptic CA2 terminals and V1aR cells in the LS. This would have a dual function of initially exciting the V1aR cells and increasing the output from CA2, possibly leading to synaptic strengthening to ultimately modulate behavior (LTP; Huang, 1998). However, if the cells in the CA2 which project to the LS V1aR cells are not V1bR cells, this suggests a more complicated circuit requiring further investigation. Overall, the results of tracing the inputs and outputs of V1aR LS cells open many more paths of inquiry into the AVP circuitry in the brain.

Having established inputs and outputs of V1aR-expressing cells in the LS, we next asked which molecularly defined LS cell types coexpress V1aR, focusing on markers previously shown to delineate LSd and LSi subregions. This approach provided a transcriptional profile of LS cell identity and neuromodulatory receptor co-expression (Dumais and Veenema, 2016; Rigney et al., 2023). We used RNAscope *in situ hybridization* to examine V1aR mRNA co-expression with several neurochemical and receptor markers. These included oxytocin receptor (*Oxtr*), neurotensin (*Nts*), vesicular glutamate transporter 2 (*Vglut2*), vesicular inhibitory amino acid transporter (*Vgat*), hypocretin receptor 1 (*Hcrtr1*), somatostatin (*Sst*), corticotropin-releasing hormone receptor 2 (*Crfr2*), and neuron-derived neurotrophic factor (*Ndnf*). By aligning V1aR expression with these molecular markers, we aimed to identify potential subpopulations and neurochemical signatures that could underlie the distinct physiological and behavioral functions of vasopressin signaling in the LS, building on prior anatomical work linking BNST projections to regionally distinct LS targets (De Vries and Buijs, 1983; Rigney et al., 2024).

We investigated *Sst* and *Nts* as common markers with some specificity for the lateral septum dorsal and intermediate subregions (Chen et al., 2024) to see if these cell types are associated with V1aR expression. Both markers had high coexpression but this differed based on location, as coexpression with V1aR was higher for both genes in the LSd compared to the LSi. This was especially true in the case of somatostatin, which suggests a potential somatostatin+ subtype for V1aR cells primarily in the LSd. Similarly, *Ndnf*, another gene prominently expressed in the LSd, was also strongly co-expressed with V1aR in the LSd with less expression in the LSi. High coexpression of V1aR with these three genes suggests that *Sst*, *Nts*, and *Ndnf* are the majority cell types that express V1aR in the LSd.

*Crfr2*, encoding the stress-modulating CRHR2 (Anthony et al., 2014) showed remarkable coexpression with V1aR and a high specificity for the LSi, suggesting its own potential region-specific cell type. Notably, this could act as a potential explanation of AVP-based anxiogenic effects if such activity results in excitation of the coexpressed *Crfr2* cell type. Taken together, *Ndnf* and *Crfr2* represent notable markers with high coexpression with V1aR for the LSd and LSi respectively.

The pattern of V1aR/Oxtr coexpression suggests that the responsiveness to AVP is weakest in the LSv and of oxytocin within the LSd, with the LSi as a region where both neuropeptides likely have conjoint activity. The highest level of V1aR expression was in the LSd, despite the fact that this region receives minimal AVP innervation (Rood et al. 2013) compared to the LSi and LSv. One possible explanation for this apparent paradox is that heightened expression of V1aR in LSd represents a need for increased sensitivity to distally released AVP, likely from hypothalamic sources (Brown et al., 2020). Another explanation is that these LSd V1aR cells may primarily be responsive to oxytocin due to the known cross-talk between the two neuropeptides and their respective receptors (Song and Albers, 2018).

## Conclusion

This validated *Avpr1a-*iCre mouse model now provides genetic access to V1aR-expressing cells, which has not been possible before, and opens the door for circuit-level functional and behavioral analyses. Similar approaches, such as the *Oxt*-Cre and *Oxtr*-Cre mouse lines, have yielded major advances in understanding oxytocin receptor function and oxytocinergic circuitry. For example, activation of oxytocin neurons in *Oxt*-Cre mice enables the social transmission of maternal behavior (Carcea et al., 2021), while circuit-mapping studies have delineated projection patterns of oxytocin neurons across the brain (Marlin et al., 2015).

Our tracing results confirm reciprocal connections between populations of V1aR-expressing cells, suggest a possible modulatory connection between LS V1aR and CA2 V1bR cells, and indicate that there exists an integrated AVP/V1aR system that incorporates much of the social-behavior mouse brain. While previous work has traced AVP fibers generally as well as those responsive to gonadal hormones, (De Vries et al., 1985, Rigney et al. 2023, Rood et al., 2013, Smith et al., 2020) this study is the first to specifically target V1aR cells in the LS which offers new lines of inquiry in AVP and social behavior. Moreover, the conserved pattern of V1aR expression and connectivity across vertebrates suggests that these circuits are derived from an ancestral vasopressinergic system that has been evolutionarily maintained to regulate social, emotional, and homeostatic functions (Donaldson and Young, 2008).

The vasopressin and oxytocin receptor systems are highly homologous and often exhibit overlapping distributions and signaling properties (Song and Albers, 2018). Our findings suggest that this overlap may extend to functional interactions within the LS. The presence of V1aR-expressing cells in close proximity to, and in some cases (i.e. the posterior region of the LSi) co-expressing, Oxtr indicates that the LS may serve as an integrative hub for both AVP and OXT signaling. Such overlap may help fine-tune social and emotional behaviors, as vasopressin and oxytocin often produce opposing or complementary effects depending on the brain region and behavioral context (Neumann and Landgraf, 2012; Caldwell and Albers, 2016). In addition, V1bR (*Avpr1b*) expression is highly concentrated in hippocampal CA2 neurons (Koshimizu et al., 2012), which provide one of the strongest monosynaptic inputs to LS V1aR cells, as confirmed by our retrograde tracing results. This connectivity suggests that any V1aR-V1bR interactions influencing LS function would most likely occur through CA2 projections onto V1aR-expressing LS cells. This aligns with prior evidence that CA2 neurons, through their vasopressin-sensitive signaling, modulate social memory and aggression and relay hippocampal information to septal and hypothalamic circuits (Leroy et al., 2018; Boyle et al., 2023). Thus, the integration of CA2-derived V1bR input with V1aR signaling in LS may represent a key pathway by which vasopressin shapes social and emotional behaviors. Together, these findings support the view that V1aR-, V1bR-, and Oxtr-expressing circuits form an interconnected modulatory network that coordinates social behavior, stress responsiveness, and affiliative processes across multiple brain regions.

