## Supplemental Figures for "Connectivity and phenotype of vasopressin 1a receptor cells in the lateral septum"

| Test | Cre+ | | Cre- | | Two-Way ANOVA |
|---|---|---|---|---|---|
| | Male | Female | Male | Female | |
| Health Assessments | | | | | |
| Vibrasse | 0.3 ± 0.21 | 1 ± 0.42 | 0.4 ± 0.27 | 1.8 ± 0.55 | Sex: $F_{(1,36)} = 7.336$, $p = 0.01$ *<br>$\eta p^2 = 0.17$, 95% CI = [0.03, 1.00]<br><br>Tukey<br>*Cre- FvM - $p = 0.04$<br>Cre+ FvM - $p = 0.16$ |
| | | | | | Genotype: $F_{(1,36)} = 1.348$, $p = 0.25$ |
| | | | | | Interaction: $F_{(1,36)} = 0.348$, $p = 0.25$ |
| Body Weight | 25.5 ± 0.40 | 20.1 ± 0.57 | 24.6 ± 0.58 | 20.9 ± 0.81 | Sex: $F_{(1,36)} = 56.164$, $p < 0.01$*<br>$\eta p^2 = 0.61$, 95% CI = [0.43, 1.00]<br><br>Tukey<br>*Cre- FvM - $p < 0.01$<br>*Cre+ FvM - $p < 0.01$ |
| | | | | | Genotype: $F_{(1,36)} = 0.007$, $p = 0.94$ |
| | | | | | Interaction: $F_{(1,36)} = 1.960$, $p = 0.17$ |
| Body Length | 9.4 ± 0.07 | 9.04 ± 0.14 | 9.48 ± 0.06 | 9.3 ± 0.10 | Sex: $F_{(1,36)} = 6.694$, $p = 0.01$*<br>$\eta p^2 = 0.16$, 95% CI = [0.02, 1.00]<br><br>Tukey<br>Cre- FvM - $p = 0.13$<br>Cre+ FvM - $p = 0.06$ |
| | | | | | Genotype: $F_{(1,36)} = 2.473$, $p = 0.12$ |
| | | | | | Interaction: $F_{(1,36)} = 0.579$, $p = 0.45$ |
| Sensorimotor Assessments | | | | | |
| Visual Placing | 1 ± 0 | 1 ± 0 | 1 ± 0 | 1 ± 0 | Sex: $F_{(1,36)} = 1$, $p = 0.32$ |
| | | | | | Genotype: $F_{(1,36)} = 1$, $p = 0.32$ |
| | | | | | Interaction: $F_{(1,36)} = 1$, $p = 0.32$ |
| Contact Placing | 1.73 ± 0.08 | 1.8 ± 0.2 | 1.87 ± 0.09 | 1.8 ± 0.2 | Sex: $F_{(1,36)} = 0.0$, $p = 1.0$ |
| | | | | | Genotype: $F_{(1,36)} = 0.187$, $p = 0.67$ |

| | | | | | |
|---|---|---|---|---|---|
| | | | | | Interaction: $F_{(1,36)}$ = 0.187, p = 0.67 |
| Limb Clasping | 8.2 ± 0.33 | 8 ± 0.37 | 7.8 ± 0.29 | 7.4 ± 0.31 | Sex: $F_{(1,36)}$ = 0.862, p = 0.36 |
| | | | | | Genotype: $F_{(1,36)}$ = 2.394, p = 0.13 |
| | | | | | Interaction: $F_{(1,36)}$ = 0.096, p = 0.76 |
| Auditory Function | 0.9 ± 0.1 | 0.9 ± 0.1 | 1 ± 0 | 0.7 ± 0.15 | Sex: $F_{(1,36)}$ = 2.077, p = 0.16 |
| | | | | | Genotype: $F_{(1,36)}$ = 0.231, p = 0.63 |
| | | | | | Interaction: $F_{(1,36)}$ = 2.077, p = 0.16 |
| Grip Strength | 123.79 ± 11.85 | 117.67 ± 9.99 | 110.80 ± 7.83 | 107.58 ± 7.58 | Sex: $F_{(1,36)}$ = 0.249, p = 0.62 |
| | | | | | Genotype: $F_{(1,36)}$ = 1.497, p = 0.23 |
| | | | | | $F_{(1,36)}$ = 0.022, p = 0.88 |
| Olfaction | 90.75 ± 15.32 | 86.23 ± 19.02 | 143.57 ± 35.92 | 132.17 ± 32.29 | Sex: $F_{(1,36)}$ = 0.087, p = 0.09 |
| | | | | | Genotype: $F_{(1,36)}$ = 3.330, p = 0.08 |
| | | | | | Interaction: $F_{(1,36)}$ = 0.016, p = 0.90 |
| Rotarod | 191.23 ± 19.20 | 228.73 ± 9.78 | 188.73 ± 23.59 | 202.97 ± 22.04 | Sex: $F_{(1,36)}$ = 1.776, p = 0.19 |
| | | | | | Genotype: $F_{(1,36)}$ = 0.530, p = 0.47 |
| | | | | | Interaction: $F_{(1,36)}$ = 0.359, p = 0.55 |
| Behavioral Assessments | | | | | |
| Elevated Plus Maze Time in Open Arm | 16.4 ± 5.94 | 19.9 ± 4.34 | 14.9 ± 3.62 | 26.4 ± 6.53 | Sex: $F_{(1,36)}$ = 2.048, p = 0.16 |
| | | | | | Genotype: $F_{(1,36)}$ = 0.228, p = 0.64 |
| | | | | | Interaction: $F_{(1,36)}$ = 0.582, p = 0.45 |
| Open Field Distance Traveled | 2759.2 ± 131.06 | 3442.4 ± 244.60 | 3197.5 ± 155.62 | 3703.2 ± 312.31 | Sex: $F_{(1,36)}$ = 7.112, p = 0.01*<br>$\eta p^2$ = 0.16, 95% CI = [0.02, 1.00]<br><br>Tukey<br>Cre- FvM - p = 0.16<br>*Cre+ FvM - p = 0.02 |
| | | | | | Genotype: $F_{(1,36)}$ = 2.458, p = 0.13 |
| | | | | | Interaction: $F_{(1,36)}$ = 0.159, p = 0.69 |

| | | | | | |
|---|---|---|---|---|---|
| Open Field Time Moving | 111.4 ± 5.45 | 141.6 ± 10.41 | 133.7 ± 6.83 | 152.7 ± 14.68 | Sex: $F_{(1,36)}$ = 6.049, $p$ = 0.02*<br>$\eta p^2$ = 0.14, 95% CI = [0.01, 1.00]<br><br>Tukey<br>Cre- FvM - $p$ = 0.26<br>*Cre+ FvM - $p$ = 0.02<br>Genotype: $F_{(1,36)}$ = 2.788, $p$ = 0.10<br>Interaction: $F_{(1,36)}$ = 0.313, $p$ = 0.58 |
| Open Field Time in Center | 96.6 ± 15.45 | 86.6 ± 3.69 | 90.3 ± 7.10 | 95.3 ± 12.44 | Sex: $F_{(1,36)}$ = 0.055, $p$ = 0.82<br>Genotype: $F_{(1,36)}$ = 0.013, $p$ = 0.91<br>Interaction: $F_{(1,36)}$ = 0.492, $p$ = 0.49 |
| Marble Burying Percent Fully Buried | 0.19 ± 0.05 | 0.16 ± 0.06 | 0.22 ± 0.05 | 0.13 ± 0.04 | Sex: $F_{(1,36)}$ = 1.754, $p$ = 0.19<br>Genotype: $F_{(1,36)}$ = 0.0, $p$ = 1.0<br>Interaction: $F_{(1,36)}$ = 0.388, $p$ = 0.54 |
| Marble Burying Percent Partially Buried | 0.39 ± 0.05 | 0.32 ± 0.05 | 0.29 ± 0.07 | 0.39 ± 0.04 | Sex: $F_{(1,36)}$ = 0.048, $p$ = 0.827<br>Genotype: $F_{(1,36)}$ = 0.146, $p$ = 0.71<br>Interaction: $F_{(1,36)}$ = 2.295, $p$ = 0.139 |
| Digging Latency | 61.3 ± 18.33 | 22.98 ± 10.35 | 32.45 ± 11.41 | 34.26 ± 11.56 | Sex: $F_{(1,36)}$ = 1.885, $p$ = 0.18<br>Genotype: $F_{(1,36)}$ = 0.437, $p$ = 0.513<br>Interaction: $F_{(1,36)}$ = 2.278, $p$ = 0.14 |
| Total Time Spent Digging | 34.22 ± 9.51 | 36.92 ± 7.00 | 38.13 ± 8.75 | 34.18 ± 7.06 | Sex: $F_{(1,36)}$ = 0.006, $p$ = 0.94<br>Genotype: $F_{(1,36)}$ = 0.005, $p$ = 0.94<br>Interaction: $F_{(1,36)}$ = 0.166, $p$ = 0.69 |
| Forced Swim Percent Time Immobile | 0.64 ± 0.04 | 0.71 ± 0.04 | 0.68 ± 0.03 | 0.65 ± 0.05 | Sex: $F_{(1,36)}$ = 0.231, $p$ = 0.63<br>Genotype: $F_{(1,36)}$ = 0.054, $p$ = 0.82<br>Interaction: $F_{(1,36)}$ = 1.715, $p$ = 0.20 |
| Sucrose Preference | 76.4 ± 4.35 | 87.89 ± 2.54 | 74.73 ± 7.51 | 87.04 ± 1.53 | Sex: $F_{(1,36)}$ = 6.703, $p$ = 0.01*<br>$\eta p^2$ = 0.16, 95% CI = [0.02, 1.00]<br><br>Tukey<br>Cre- FvM - $p$ = 0.13 |

| | | | | | |
|---|---|---|---|---|---|
| | | | | | *Cre+ FvM - p = 0.04 |
| | | | | | Genotype: $F_{(1,36)}$ = 0.078, p = 0.78 |
| | | | | | Interaction: $F_{(1,36)}$ = 0.009 , p = 0.93 |
| Sucrose Test - Total Fluid Consumption | 8.5 ± 2.05 | 2.1 ± 0.43 | 7.2 ± 1.46 | 2.4 ± 0.27 | Sex: $F_{(1,36)}$ = 5.222, p = 0.03*<br>$\eta p^2$ = 0.13, 95% CI = [0.01, 1.00]<br><br>Tukey<br>*Cre- FvM - p = 0.02<br>Cre+ FvM - p = 0.30 |
| | | | | | Genotype: $F_{(1,36)}$ = 0.001, p = 0.97 |
| | | | | | Interaction: $F_{(1,36)}$ = 0.31, p = 0.58 |

**Table 1**. Results of health, physiological, and behavioral assessments. 2-Way ANOVA with Tukey’s HSD post-hoc tests. *p<0.05

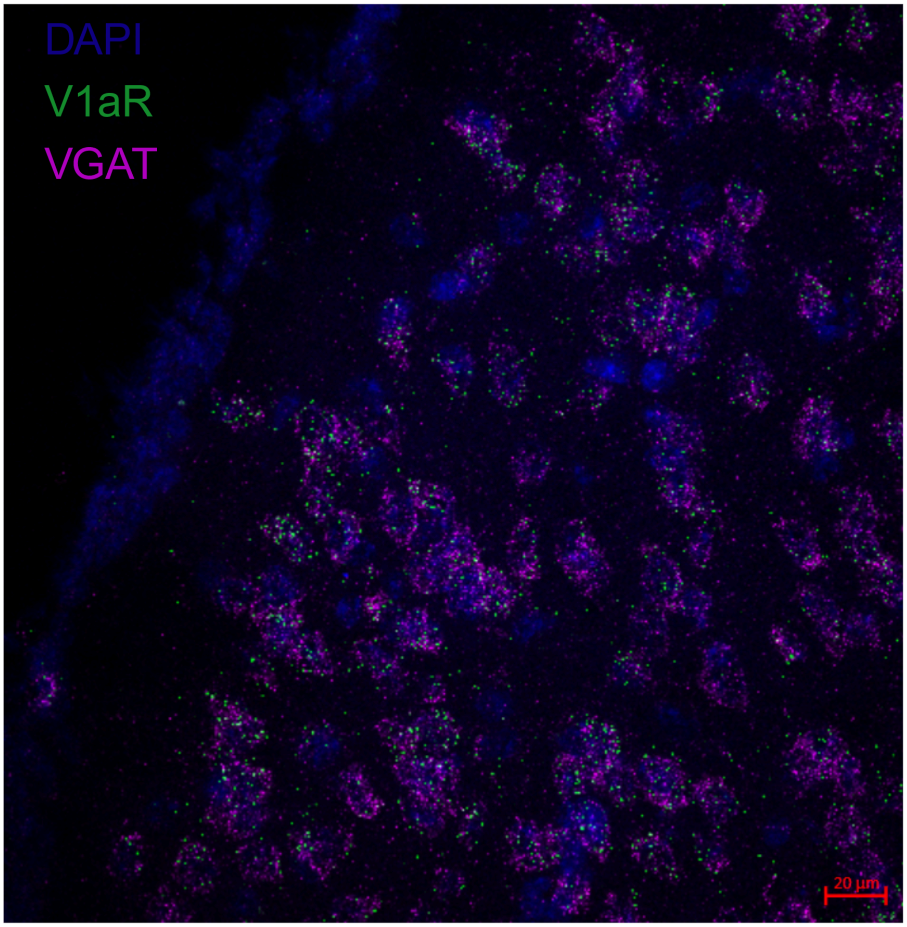


**Supplemental Figure 1.** *In situ* hybridization showing V1aR mRNA (green), and VGAT mRNA (magenta) colocalization in the LS.
